# Global motor dynamics - Invariant neural representations of motor behavior in distributed brain-wide recordings

**DOI:** 10.1101/2023.07.07.548122

**Authors:** Maarten C. Ottenhoff, Maxime Verwoert, Sophocles Goulis, Louis Wagner, Johannes P. van Dijk, Pieter L. Kubben, Christian Herff

## Abstract

Neural activity is presumed to be correlated with motor behavior almost everywhere in the brain, implying that many different brain processes are involved in generating a behavioral output. Recent studies in multiple non-human species have observed pervasive brain-wide neural activity directly related to motor output. However, similar brain-wide investigations in humans with high-temporal resolution are lacking to date. Here, we recorded invasive data from brain-wide distributed electrodes in humans and reveal global neural dynamics that are predictive of movement across tasks and across participants. The dynamics are remarkably stable between participants with substantially varying electrode configurations and to loss of information. We demonstrate that these global neural dynamics are near brain-wide and present in all participants. Uncovering these global neural dynamics may allow for a more holistic and network-based perspective on motor-related neural activity.

## 1 Introduction

Nearly all decisions eventually lead to movement, and consequently most neural processes facilitate a downstream motor output. While the neural basis and localization of the processes involved remain elusive, it is conceivable that movement sparks the involvement of many different brain areas. In animals, brain-wide neural activity associated to motor output was observed in zebrafish^1^ and mice^2–5^, where spontaneous behaviors elicited a broadcast of neural activity throughout the dorsal cortex. Moreover, in a study with 12.000 recorded neurons in mice, nearly all neurons were strongly modulated by any type (instructed and uninstructed) of movement in mice^6^. Consequently, the authors conclude that robust movement representations might be present throughout the brain. In humans, early indications in functional magnetic resonance imaging (fMRI) suggest that similar brain-wide activity exists^7^, but it has yet to be demonstrated in humans with high temporal resolution in electrophysiological recordings.

Capturing the neural substrate of these brain-wide motor-related responses in humans will not only increase our understanding of motor processes in the brain, but also allow for robust neural decoders. Results from animal studies that identified global motor-related activity suggest that there indeed exists decodable information throughout the brain. To this end, stereotactic-electroencephalographic (sEEG), also called depth electrodes, provide a unique opportunity to record from sparse but brain-wide cortical and subcortical areas, and allow us to shine light on global motor activity and its dynamics^8^. To date, motor decoding studies based on sEEG recordings have described the decoding of action (e.g. different hand signs or grasp types) from a variety of individual cortical and subcortical areas^9^, including the ventral premotor cortex^10^, posterior parietal cortex^11–13^, somatosensory cortex^10^, supramarginal gyrus^10,12,14^, middle temporal gyrus and fusiform gyrus^14^, insula^12,14^ and hippocampus^12,14^.

Given that individual areas are predictive of movements, the underlying neural dynamics between areas could increase the available information and uncover global dynamics. To capture these dynamics into a low-dimensional representation, multiple techniques are available to reduce the neural space into a single manifold that describe the underlying neural dynamics^15–20^. Manifolds are demonstrated to capture robust representations of latent dynamics^21^ and to be predictive for multiple types of behaviors^22,23^and subjects^24^. In the context of decoding motor behavior, these neural manifold approaches have almost exclusively been used to decode spiking activity recorded in the primary motor cortex, although the methods are well suited for local field potentials recorded with sEEG electrodes.

Here, we explore motor-related activity from brain-wide distributed recordings. Eight participants implanted with sEEG electrodes performed trial-based executed and imagined motor tasks. In each 3-second trial, they continuously opened and closed either their left or right hand. In total, we recorded data from 956 contacts covering 60 unique brain areas. By extracting a low-dimensional representation, we were able to decode movements across tasks and participants using a Riemannian geometry approach (Fig. 1, Fig. 2a). We find remarkably stable motor dynamics that are independent of task and participant. Indeed, we were able to decode movements across participants with non-overlapping electrodes.

**Figure 1.**
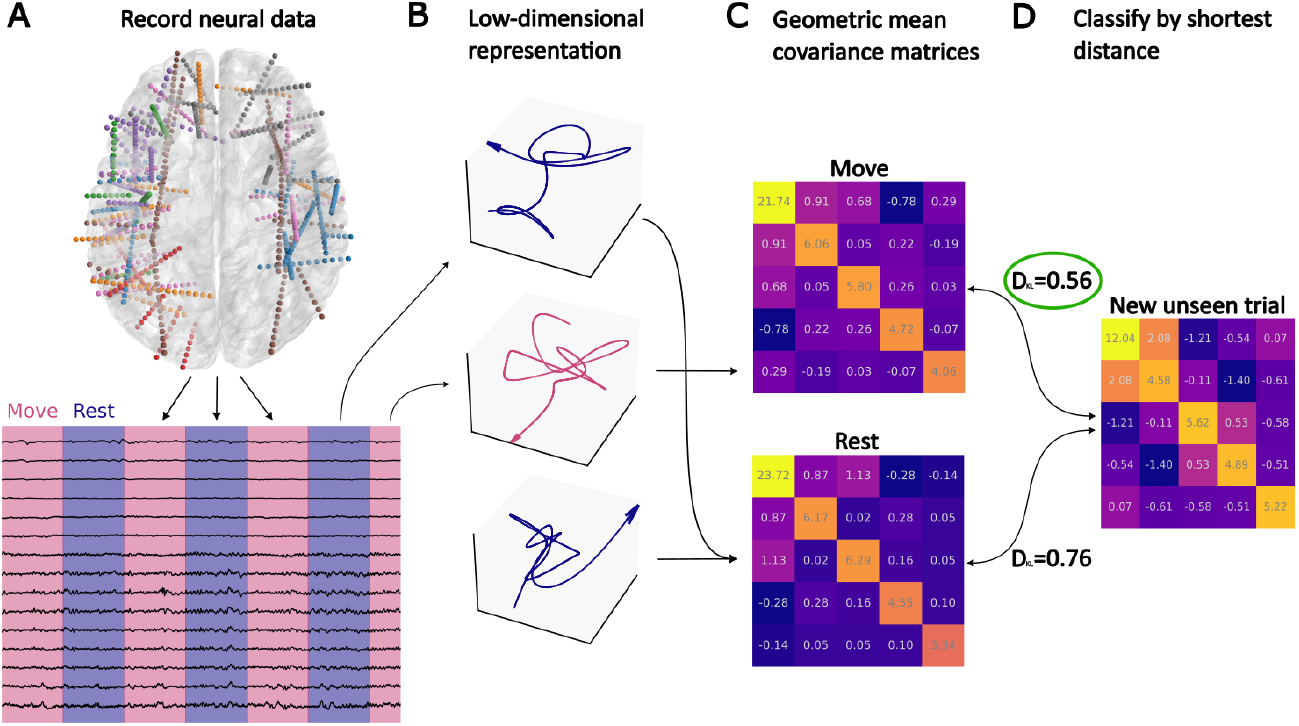
Conceptual overview of the methods. **A** - Electrode configurations of 8 participants mapped onto an average brain. A total of 956 contacts cover 60 unique brain areas. For each participant, data were recorded in 3-second trials in either a move (red) or rest (blue) trial. **B** - On the training set, principal components were extracted and each trial was transformed to a lower dimensional representation. **C** - Then, the sample covariance matrix was estimated for each trial, and for all trials per class in the training set, the geometric mean of the covariance matrices was calculated using the kullback-leibler divergence. **D** - New, unseen trials from the test set are then classified by findings the geometric class mean with the smallest kullback-leibler divergence to the new trial sample covariance matrix.

**Figure 2.**
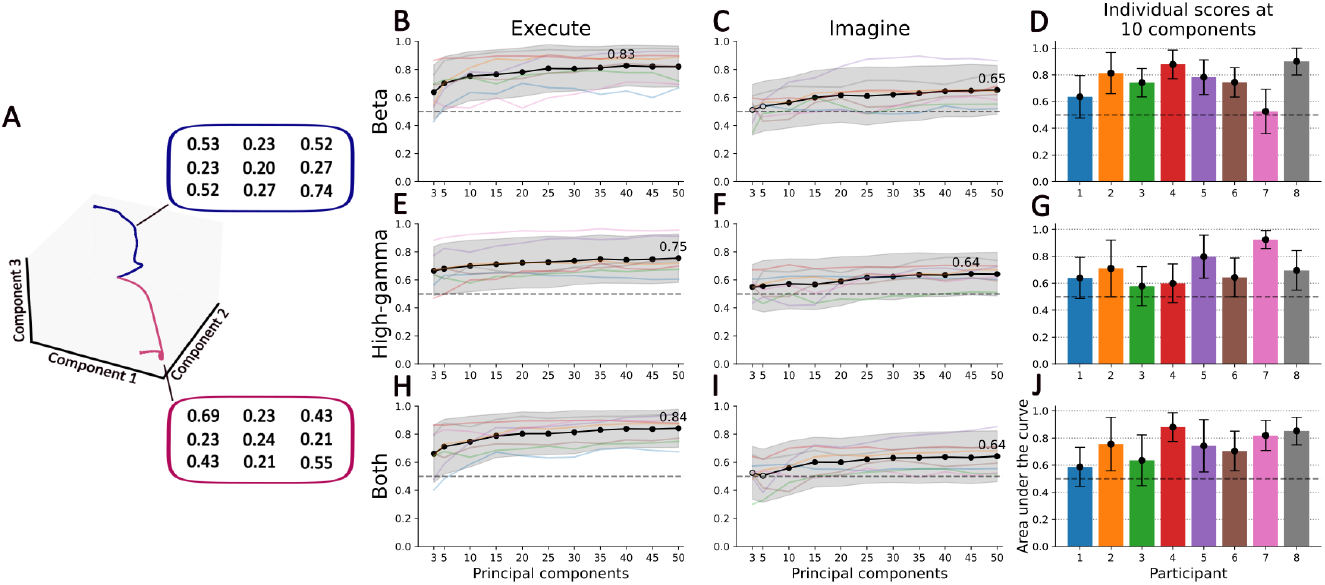
Decoder performance. **A** - Representation of the average dynamics of movements and rest in the first three components. Blue (rest) and red (move) boxes illustrate the corresponding covariance matrices. **B** - Average decoding performance for executed movements using only beta power. Performance significantly (one sample t-test, *α* = 0.05, false discovery rate corrected) above chance (0.5, black dotted line) is shown by filled circled, non-significant scores are shown as open circles. The annotated number shows the maximum performance. Each colored line is one participant. **C** - Same as **B**, but for imagined movement decoding. **D** - Decoding performance per participant for executed movements using only beta power and 10 principal components. Error bars show standard deviation over folds. Red dotted line shows chance level (0.5). **E, F, G** - Same as **B, C, D**, respectively, but with high-gamma power. **H, I, J** - Same as **B, C, D**, respectively, but with both beta and high-gamma power.

## 2 Results

### 2.1 Executed and imagined movement can be decoded from low-dimensional neural representations from distributed brain areas

The Riemannian geometry-based classifier was able to decode executed movements for beta, high-gamma and both frequency bands and all number of principal components (Fig. 2b, e, h) significantly above chance level. Beta power resulted in higher decoding performance than high-gamma power (0.83 *±* 0.15 area under the curve (auc) vs 0.75 *±* 0.17 auc), and including both frequency bands led to similar performance (0.84 *±* 0.14 auc) as beta power. All number of principal components and frequency inputs resulted in decoding performance significantly above chance (one sample t-test, *α* = 0.05, FDR corrected). Zooming in on 10 components for executed movements, based on the optimum further described in section 2.2, most participants reached above chance decoding, averaging 0.75±0.18, 0.69 *±* 0.18, 0.74 *±* 0.18 for beta, high-gamma and both, respectively (Fig. 2d, g, j). Inter-participant variance was high, ranging from chance level decoding to *auc >* 0.9. This is regularly observed in sEEG decoding studies due to varying electrode configurations. In the imagined task, the decoder reached above chance decoding as well for nearly all combinations of principal components and frequency bands (Fig. 2c, f, i), except for the lower number of principal components (*n*_*components*_ *<*= 10). Compared to executed movement, the performance was lower overall, but similar between all frequency bands (0.65 *±* 0.17, 0.64 *±* 0.15 and 0.64 *±* 0.18 auc), where beta power and beta & high-gamma power achieved the highest performance. Given that beta provides the same predictive power as both beta and high-gamma, further analyses will continue with beta power only.

### 2.2 The optimal amount of components remains stable under loss of information

To assess whether the extracted low-dimensional representation captures neural dynamics on a stable manifold, we performed an ablation study that progressively removed information from randomly selected channels. The number of principal components included strongly determined the baseline performance (Fig. 3a), defined as having access to 100% of the available channels. A higher baseline performance led to a progressive decrease in stability, as the performance dropped more rapidly for models with more components when information was removed. Using *n*_*components*_ *<* 10, the performance showed no drop in performance for up to 50% information loss, whereas *n*_*components*_ = 50 led to a drop of 0.15 auc when losing only 10% of information. Losing even more information reduced the performance to chance level already at 70% of the available channels. The results introduce a trade-off between decoder performance and stability, suggesting that there exists an optimum. Therefore, we calculated the mean performance over all percentages of information loss for each number of principal components, and find a smooth curve with an optimum of *n*_*components*_ = 10 (Fig. 3b). Further inspection of the individual scores at the optimum (Fig. 3c) revealed that the stability found on average holds true for each participant, regardless of their individual performance. While standard deviation increased slightly as expected, the mean performance remains equal or decreases only slightly under increasing loss of information.

**Figure 3.**
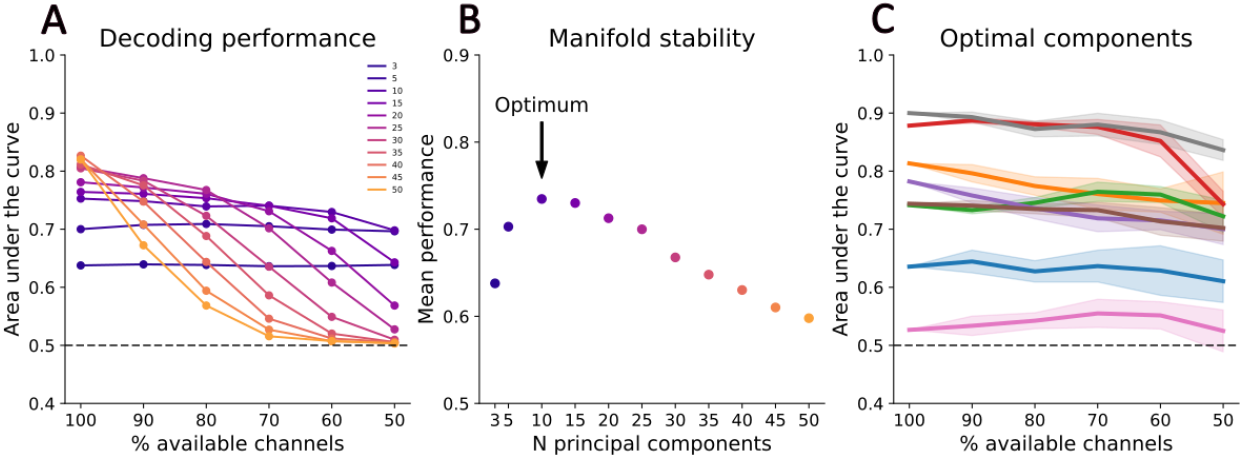
Manifold stability. **A** - Decoder performance when information is removed from a percentage of channels. Less components (purple) leads to lower baseline performance that remains stable over information loss, whereas more components (yellow) leads to higher baseline performance, but instability to information loss. **B** - Mean performance over different percentages of information loss, where the optimum is shown at 10 principal components. **C** - Performance per participant for the optimal number of components. Regardless of baseline performance, the performance remains relatively stable with increasing information loss for each participant.

### 2.3 Optimal manifold captures similar information across tasks

Next, we explored whether the stable manifold describes task-specific or broader movement-related information. To test this, we trained our decoder on executed movements and tested it on imagined movements. The decoder reached performance significantly above chance (one sample t-test, *α* = 0.05, FDR corrected) between 10 to 25 principal components and 40 components (Fig. 4a, black line). Maximum performance was 0.61 *±* 0.07 auc at 25 components. Interestingly, cross-task performance is equal to within-task performance (Fig. 4a, black dashed line), as there is no significant difference between the performances. The performance is similar specifically up to 30, after which both start to diverge with an increasing number of principal components.

**Figure 4.**
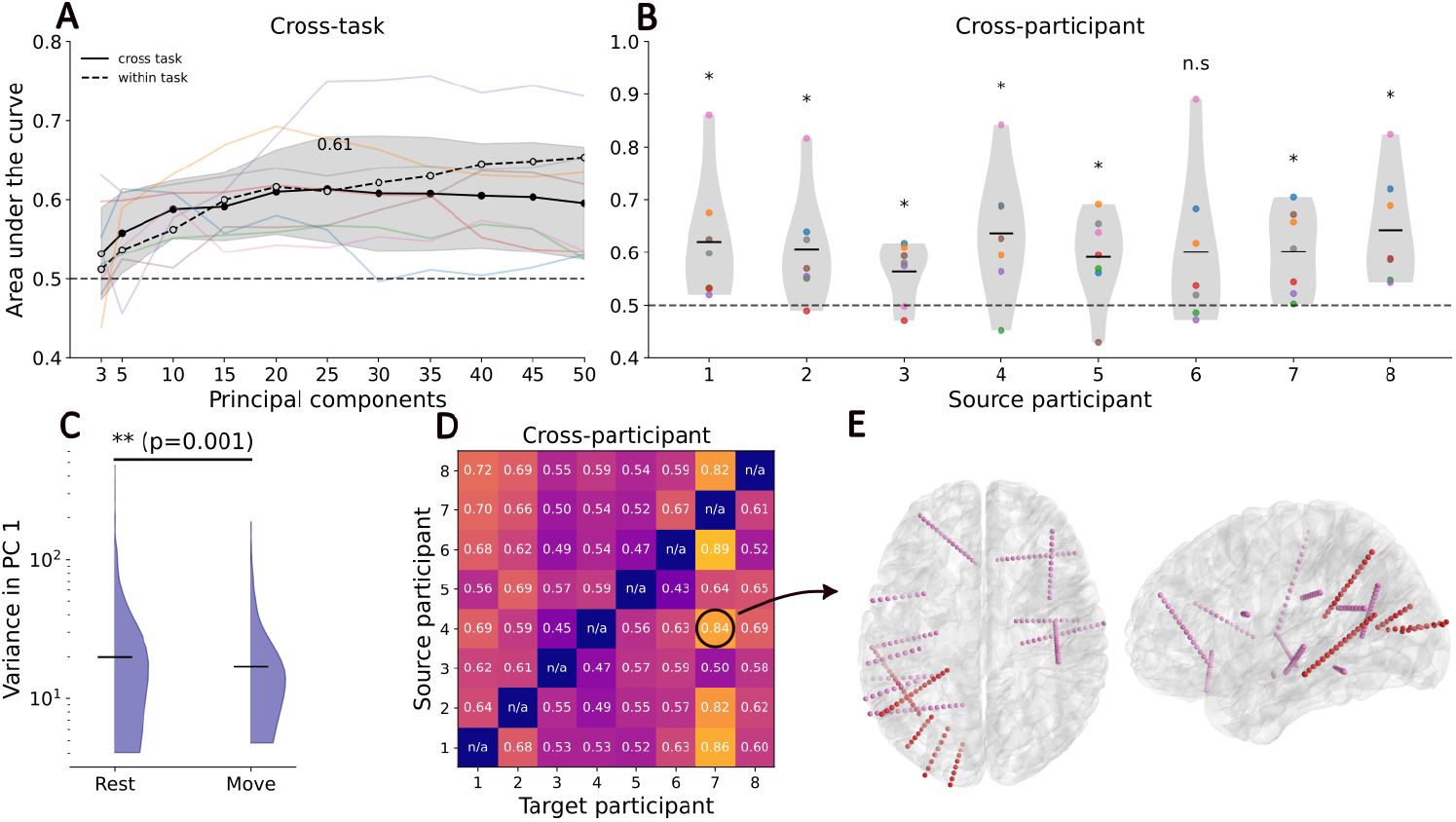
Cross-task and cross-participant decoding. **A** - Performance across task, where the decoder is trained on executed movements and tested on imagined movements. For cross-task (filled line) the filled circles represent scores significantly above chance (one sample t-test, *α* = 0.05, FDR corrected), whereas the circles in within tasks (dashed line) shows whether the scores are significantly different then across task (independent t-test, *α* = 0.05, FDR corrected). None of the within task decoding scores are significantly different than across task, although performance starts to increase gradually over cross-task performance with 30 or more principal components. **B** - Decoding performance when trained on a source participant and tested on the remaining target participants. Each color represents a single participant. * shows decoding performance significantly above chance (one sample t-test, *α* = 0.05, FDR corrected). n.s. = not significant. **C** - Distribution of variance in the first components per class. PC = principal component. **D** - Performance matrix for each source-target pair. An example performance is highlighted by the circle and arrow, and shows the **E** - electrode configurations of the source (p4, red) and target (p7, pink) participant.

**Figure 5.**
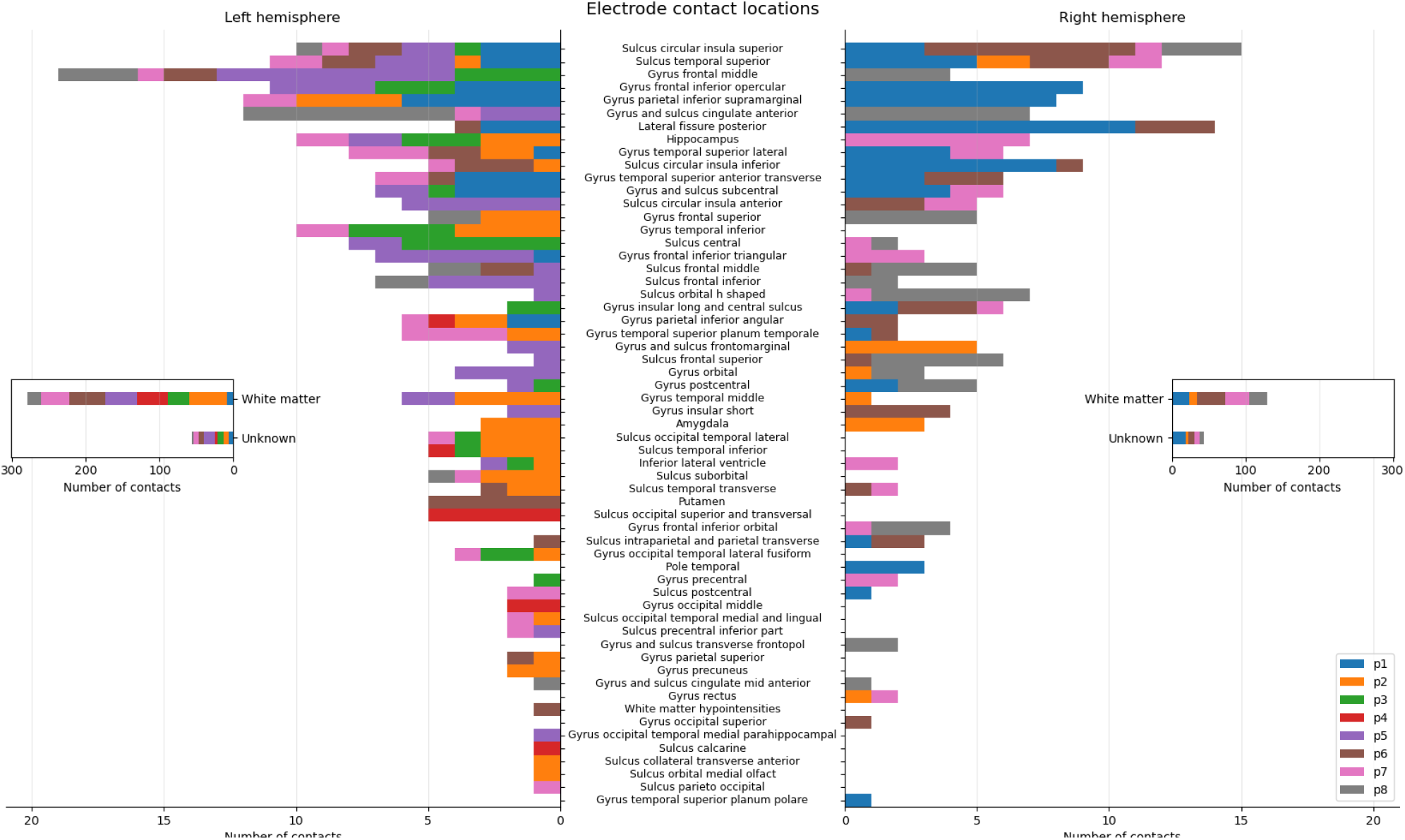
Brain regions covered by the implanted electrodes. Each color is a different participant. Note that most contacts are in white matter and unknown areas, as shown in the insets.

### 2.4 Optimal manifold captures participant invariant global neural dynamics

To further explore the extend of shared information captured by the manifold, we evaluated the decoding performance on executed movements across participants. We trained our decoder on a source participant and tested it on each remaining target participant. For 7 out of 8 source participants the decoder was able to decode significantly above chance (one sided t-test, *α* = 0.05, FDR corrected, Fig. 4b) on average over all target participants using beta power. For the same analysis using high-gamma power, 6 out of 8 participants reached significantly above chance decoding, whereas beta + high-gamma resulted significant decoding in 0 out of 8 participants. While each participant’s electrode configuration was quite varied and distributed throughout the brain (Fig. 1b, Fig. 4e), nearly all electrode configurations were sufficient to capture similar information across participants on average. Surprisingly, even source-target pairs with barely overlapping electrode configurations were able to achieve good decoding results (Fig. 4d, e). A closer inspection of the first principal component showed that the variance is significantly lower (independent t-test, *p* = 0.001) during the move trials compared to rest (Fig. 4c). This suggests that the first component captures beta band desynchronization, as this analysis only uses beta power as input. Inspecting the individual performance values showed that the source-target pairs are not symmetrical: if one participant is a good source, it does not seem to predict that one is a good target as well. For example, the circle shown in Fig. 4d, shows a high decoding performance as target (*auc* = 0.82), but the reversed direction only shows *auc* = 0.54. Furthermore, it seems that if a participant has good target decoding, they are generally a good target for the other participants (Fig. 4d, columns). From a source perspective, this is not the case, as the scores vary substantially over participants. Quantitatively, it shows that the standard deviation (std) of the target perspective, that is whether a participant is a good target, is substantially lower than the source perspective: 0.062 *±* 0.026 vs 0.096 *±* 0.020 (mean ± std of the std). Taken together, our results suggests that decodable global neural dynamics exist and these dynamics are similar across tasks and across participants.

## 3 Discussion

We find a stable low-dimensional manifold underlying global motor activity in sEEG recordings that is predictive for both executed and imagined grasping movements. Any number of components is sufficient to decode executed movement, but we observe that including more components gradually increases performance (Fig. 2b, e, h). In imagined movements, the decoding performance is significantly above chance from 15 or more components, and the same gradual increase in performance is observed (Fig. 2c, f, i). However, the ablation study revealed that including too many components decreased the stability of the decoder. The more components included, the faster the performance of the decoder decreased under loss of information (Fig. 3). Thus, it seems like restricting the number of components increases the generalizability of the decoder and prevents it from fitting dataset specific information. It is challenging to discern whether the increased performance is movement-related information only captured by the participant-specific electrode configuration, or that it increasingly overfits on dataset specific bias or other non-movement related noise.

Our results indicate that there is a performance-stability trade-off (Fig. 3b) with an optimum of 10 components. Decoding with this optimum reveals a manifold that describes remarkably stable neural dynamics, demonstrating that movement-related activity is captured in a smaller subspace than the original space. For all participants, this manifold remained stable for up to 50% of missing information, regardless of baseline performance. The stability shows that the information must come from multiple sources, suggesting that the manifold captured a distributed network throughout the brain. If a single region or process was mainly driving performance, we would expect that fewer components were required to capture the same information, a larger standard deviation under loss of information, as well as non-significant decoding performance across participants. The distributed network captured by the manifold is even predictive across tasks and across participants with non-overlapping electrode configurations (Fig. 4a, b), demonstrating that it describes generalizable dynamics. In the cross-task analysis (Fig. 4a) the decrease in generalizability becomes apparent when adding more components. For up to 25 components, the performance is equal, after which the within-task performance continues to increase and cross-task performance starts to decrease.

Taken together, our results point towards global motor dynamics and are congruent with results reported in other species^2–4^, other recording modalities^7^, and in human sEEG motor decoding studies decoding from individual brain-wide areas^9–14^. A key difference with the global motor dynamics reported in animals or fMRI is that those are based on spiking activity or hemodynamics, while our results are based on local field potentials. Furthermore, Steinmetz et al.^2^ report that a majority of the neurons correlated with movement increased their activity, while a significant minority reduced their activity, highlighting the heterogeneity in responses required to capture the full neural dynamics. In this work, we focused on beta and high-gamma activity because of their known involvement with motor behavior. We found a significant decrease in variation in the first components between move and rest that might indicate a beta desynchronization (Fig. 4c). Nonetheless, perhaps a broader spectral scope could reveal stronger generalizable dynamics.

Our data reveal some contours of the network. First, the network seems to be global, but not ubiquitous: not all participants reached sufficient decoding performance when using the optimal number of components, (Fig. 2d, g, j) and some participants seems to be good targets but poor sources for decoding, and vice versa (Fig. 4b, d). Notably, we observe that the lowest performing participant in executed beta (p7, pink) is the best target for cross-decoding (Fig. 4). Even more, based on the individual source-target scores, some participants are good targets for all other participants. That means that if a participant is a good target in a source-target pair, then it is likely that the participant is also a good target for the other participants. However, this does not seems to be the case from the source perspective. The varying performance between participants and source-target pairs might also be influenced by non-technical reasons, such as varying engagement, as all our participants are under clinical treatment during our experiments, and often report tiredness and lack of concentration.

The global motor dynamics may enable us to combine data from multiple participants, regardless of electrode configuration, improving performance of future decoders as well as reducing calibration times. Although the task is simple, detecting movements is an essential part for hierarchical decoders^25^,^26^. Moreover, movement detection might be useful for adaptive deep-brain stimulation, where an intended movement might be detected, which subsequently activates stimulation^27^. More speculatively, the revealed global neural dynamics might be able to inform about a disease state^28^, where changes in global motor dynamics might reflect disease progression. Future work will be required to disentangle the size and content of the neural dynamics, and explore potential application in new or existing decoders.

## 4 Conclusion

In summary, we have identified decodable global motor-related neural dynamics that is captured by a low-dimensional manifold. This manifold is stable to loss of information, and captures information that is similar across tasks and across participants, even with varying and non-overlapping electrodes. These results are the first demonstration showing decodable brain-wide movement-related neural activity in human electrophysiological recordings, and builds upon studies showing similar brain-wide activity across multiple species. These global dynamics might open up the way for a broader scope for all movement-related neuroscience research, including combining datasets of multiple participants, detection of movements for adaptive stimulation technologies and potentially disease progressions states.

## 5 Methods

### 5.1 Participants

We recorded data from eight epilepsy patients (age 35.8 *±* 14.2, mean *±* standard deviation, supplementary table 1). All participants were undergoing presurgical assessment for resection surgery as treatment for their medication-resistant epilepsy. Each participant was implanted with a varying amount of sEEG electrodes (supplementary table 1) in varying locations (Fig. 1a, supplementary Fig. 5). All electrode configurations and trajectories were solely based on clinical need and were not influenced in any way by this study. The amount of implanted shafts ranged from 5 to 14 electrodes, containing 42 to 127 recordable contacts.

**Table 1.**
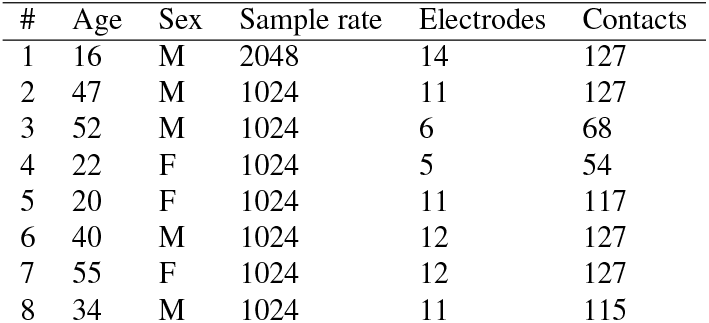
Participant information. Age = years, Sample rate = Hz

| # | Age | Sex | Sample rate | Electrodes | Contacts |
| --- | --- | --- | --- | --- | --- |
| 1 | 16 | M | 2048 | 14 | 127 |
| 2 | 47 | M | 1024 | 11 | 127 |
| 3 | 52 | M | 1024 | 6 | 68 |
| 4 | 22 | F | 1024 | 5 | 54 |
| 5 | 20 | F | 1024 | 11 | 117 |
| 6 | 40 | M | 1024 | 12 | 127 |
| 7 | 55 | F | 1024 | 12 | 127 |
| 8 | 34 | M | 1024 | 11 | 115 |

### 5.2 Ethical approval

The institutional review board of Maastricht University and Epilepsy Center Kempenhaeghe (METC 20180451) approved the experimental protocol. All experiments were in accordance with local guidelines and regulations, and were under supervision of experienced healthcare staff. All participants participated voluntarily and provided written informed consent.

### 5.3 Tasks

Our participants performed an executed and imagined continuous grasping task. In the execution task, they were instructed to open and close their left or right hand continuously, based on visual instruction. Each trial was 3 seconds long, directly followed by a 3 second rest period (Fig. 1a). Left and right hand were each cued 30 times in pseudorandomized order, resulting in a total of 60 movement trials and 60 rest trials. After a short break, the participants were instructed to imagine performing the previous execution task. The imagined task was always after the execution task. In earlier pilots, participants reported to find it challenging to imagine the movements. Thus, performing the executed task prior to the imagined task provided a fresh memory of the proprioceptive and kinematic experience of the grasping movement. To further aid the participants, the experimenter briefly introduced a kinesthetic and visual strategy^29^. However, the participants were free to use any strategy they felt was most effective. During the imagery task, the experimenter instructed the participants to remain completely still and visually checked whether they adhered to the instruction.

### 5.4 Recordings and electrodes

Participants were implanted with platinum-iridium stereotactic-electroencephalography electrodes (Microdeep intracerebal electrodes; Dixi Medical, Beçanson, France), containing 5 to 18 contacts (2 mm long, 0.8 mm in diameter and 1.5 mm intercontact distance). Neural activity was recorded using two stacked Micromed SD LTM amplifiers (Micromed, S.p.A., Treviso, Italy). All contacts were referenced to a contact in white matter that visually did not show epileptic activity, determined by the epileptologist. Neural data and experimental stimuli were synchronized using LabStreamingLayer^30^. In this work, we refer to electrodes as the full shaft and contact as each recording location on the shaft. Once data is recorded and digitized, we then refer each contact as a channel.

### 5.5 Imaging

We determined the anatomical locations of each contact by coregistering a pre-implantation anatomical T1-weighted MRI scan and post-implantation CT scan. We first parcellated the MRI using Freesurfer (https://surfer.nmr.mgh.harvard.edu/) and then labeled the anatomical locations according to the Destrieux atlas^31^ using img_pipe^32^. To generate a visualization with all electrodes from all participants on a single brain (Fig. 1a), we warped all brains and corresponding electrode locations to the CVS average-35 atlas in MNI512 space. Note that anatomical locations are always determined in native space.

### 5.6 Electrode coverage

A total of 956 contacts on 82 electrodes covered 59 unique grey matter areas (supplementary Fig. 5). The grey matter areas covered most were the insular sulcus (*n* = 25), superior temporal sulcus (*n* = 23) and middle frontal gyrus (*n* = 23). Most non-grey matter areas were in white matter (*n* = 408) or unknown areas (*n* = 100). The unknown areas could not be labeled due to various reasons, such as the atlas not having a label for a specific location or contacts between brain tissues, e.g. in sulci. technical reasons.

### 5.7 Data preparation

All our analyses were done using Python 3.9.7 and all code is openly available on (TODO github) First, we removed channels without relevant information, such as marker channels and disconnected channels. Then we removed the channel mean and detrended the signal. Lastly, we extracted 3 sets of frequency bands: beta (12 *−*30 hz), high-gamma (55 *−*90 Hz) and both. For the combination of both frequency bands, we concatenated the channels per frequency. To acquire the instantaneous power, we band-pass filtered the data using a zero-phase finite impulse response filter and then took the absolute of the hilbert transform. Finally, we split the continuous data into trials, and combine left and right hand movement trials into a single move class.

### 5.8 Decoding

We used a Riemannian classifier, which directly classifies based on the trial covariance matrix^33^. Riemannian approaches have shown to be promising for brain-computer interface applications, given its robustness to outliers and applications in surface EEG^34,35^. To decode movement from neural activity, we first split the data into training and test data using 10-fold cross validation. On the training data, we z-score the data, fit a principal components analysis (PCA) and transform the data to 3 to 50 principal components (Fig. 1b). Then, for each trial we estimate a regularized (Ledoit and Wolf^36^) sample covariance matrix and calculate the geometric mean of all covariance matrices of one class using the symmetric Kullback-Leibler divergence^37^ (Fig. 1c). To classify trials in the test set, the learned standardization and filters from the PCA are applied to the test data. Then, for each trial in the test set the same regularized sample covariance matrix is estimated. Next, a prediction is made per trial by the minimum distance to the learned geometric class means (Fig. 1d). We used the pyRiemann package^38^ for covariance estimation and decoding.

### 5.9 Manifold stability

To assess whether the extracted low-dimensional representation captures neural dynamics on a stable manifold, we performed an ablation study that removed information from randomly selected channels^39^. We progressively removed information by setting all values in randomly selected channels to zero, and then retested our decoder. Baseline performance was defined at 100% of the available channels. We then removed information from 10%, 20%, 30%, 40% and 50% randomly selected channels by setting all values to 0. We then applied the previously described decoding (section 5.8) strategy. Within each fold, we repeated the random channel selection 10 times.

### 5.10 Cross-task and cross-participant decoding

To decode across tasks, we trained our decoder on the full executed dataset, and tested it on the full imagined dataset. The standardization, principal components filters and geometric class means were fitted on the executed data, and then applied to the imagined test data. To decode across participants, the same strategy was used as cross-task decoding, but the training and test set were a source and a target participant.

### 5.11 Evaluation

All decoding scores were evaluated by the area under curve the receiver operator characteristic. To compare the decoding results against chance level (*AUC* = 0.5), we used a one-sample t-test and applied a False Discovery Rate (Benjamin-Hochberg) correction to correct for multiple testing in the main results (Fig. 2b, c, e, f, h, i) and Fig. 4a, b).

